# Unveiling the Structural Proteome of an Alzheimer’s Disease Rat Brain Model

**DOI:** 10.1101/2024.07.11.602936

**Authors:** Elnaz Khalili Samani, S.M. Naimul Hasan, Matthew Waas, Alexander F. A. Keszei, Xiaoxiao Xu, Mahtab Heydari, Mary Elizabeth Hill, JoAnne McLaurin, Thomas Kislinger, Mohammad T. Mazhab-Jafari

## Abstract

Studying native protein structures at near-atomic resolution in crowded environment presents a challenge. Consequently, understanding the structural intricacies of proteins within pathologically affected tissues often relies on mass spectrometry and proteomic analysis. In this study, we utilized electron cryomicroscopy (cryo-EM) and a specific method of analysis called Build and Retrieve (BaR) to investigate structural characteristics of protein complexes such as post-translational modification, active site occupancy, and arrested conformational state in Alzheimer’s Disease (AD) using brain lysate from a rat model (TgF344-AD) of the disease. Our findings reveal novel insights into the architecture of these complexes, which we corroborate through mass spectrometry analysis. Interestingly, it has been shown that the dysfunction of these protein complexes extends beyond AD, implicating them in cancer, as well as other neurodegenerative disorders such as Parkinson’s disease, Huntington’s disease, and Schizophrenia. By elucidating the structural details of these complexes, our work not only enhances our understanding of disease pathology but also suggests new avenues for future approaches in therapeutic intervention.

## Introduction

Alzheimer’s disease (AD) is a prevalent contributor to dementia, serving as a principal factor in the increased morbidity and mortality associated with aging^1^. Despite significant advancements in comprehending the pathogenesis of AD and refining our conceptualization of the disease, the absence of disease-modifying treatments persists^2^. Recent studies showed that approximately 30% of current drug trials for AD are directed towards the canonical targets, amyloid and tau^3^. Research on AD has reached a pivotal point, as we face the fact that over 30 years of experimental work has provided limited approaches in AD treatment, with a shift in focus from traditional amyloid plaques and tau tangles to novel targets such as intracellular proteins and biochemical pathways in the realms of biomarker and drug exploration. For instance, the adverse impact of chronic inflammation on neuronal death might be addressed by developing novel anti-inflammatory agents^4^.

Characterization of AD pathobiology at different stages of disease development is challenging. As rats are 4 –5 million years closer to humans than mice in evolution^5^, therefore, Cohen et al. developed a TgF344-AD rat model, characterized by the overexpression of mutant human amyloid precursor protein (APPsw) and presenilin 1 (PS1ΔE9) genes, mirrors the independent causative factors associated with early-onset familial AD. These rats exhibit age-dependent cerebral amyloidosis, which precedes the onset of tauopathy, gliosis, apoptotic loss of neurons in the cerebral cortex and hippocampus, and cognitive disturbances^6^. In 2022, Bernaud et al. studied the behavioral cognitive decline on the female model of TgF344-AD rats and showed the impairment begins at early stages after six months^7^. Therefore, this model of AD has been widely used in research with the aim of developing new treatments^8,9^. Considering the purpose of introducing new biomarkers for AD using this rat model, proteomics and structural studies of the intracellular disease associated proteins in the brain could open new avenues for AD drug development.

By the introduction of new direct electron detectors, single-particle cryo-electron microscopy (cryo-EM) has revolutionized the determination of three-dimensional (3D) structures of biomacromolecules at near-atomic resolution^10^. This technology provides detailed insights into previously unknown structures and elucidates the mechanisms of crucial biomacromolecules smaller than 100 kDa, which is valuable for drug discovery^11^. However, challenges persist, as historical requirements for homogenous and pure samples can hinder drug design. Here, we utilized Build and Retrieve (BaR) which is an iterative methodology that can carry out in silico purification and sorting of protein complexes within a large, diverse dataset from a heterogenous sample^12^. Deploying this method in image analysis has the capability to deconvolute images, facilitating the simultaneous generation of near-atomic resolution cryo-EM maps for individual proteins within a heterogeneous, multiprotein sample^12–15^.

To exemplify the ability of BaR to elucidate the proteome of AD rat brain, we studied the near-atomic resolution structure of cytoplasmic proteins involved in axonal development, neurotransmitter recycling, cell migration and synaptic plasticity, isolated relative to their sizes from TgF344-AD rat brain. Importantly, all these enzymes are linked to the pathobiology of Alzheimer’s disease. For instance, hyperphosphorylation of the encoded dihydropyrimidinase-like 2 (DPYSL-2) protein may play a key role in the development of AD^16,17^, glutamine metabolism by the glutamine synthetase is compromised in AD patients^18^, detoxification/repair property of actin filament is increased in patients showing resilience to AD^19^ and vacuolar ATPase (V-ATPase) is a hub that mediates proteopathy of oligomeric amyloid beta peptide (Aβ) and hyperphosphorylated Tau (p-Tau)^20^. Therefore, having insight into these proteins’ structure from disease tissue directly can yield interesting points for further investigation as to the underlying pathogenesis of AD.

## Results

### 1) Protein extraction and MS validation

Following lysis of the perfused whole brain from 22 months old TgF344-AD rat brain, we proceeded to enrich proteins from the cytosolic fraction through size exclusion chromatography (Figure S1-A). This method yielded two prominent peaks in the chromatogram corresponding to molecular weights of approximately 500-600 kDa and 150-450 kDa (Figure S1-B). In the next step, a proteomic analysis was conducted on each peak to uncover the proteins comprising them. The topmost abundant proteins within these enriched fractions were determined by their peptide-spectrum matches (PSMs) (Figure S1-D). Subsequently, individual single-particle images of the two distinct enriched samples were acquired separately using cryo-electron microscopy (cryo-EM). Collected movies were subjected to data processing utilizing the BaR method (Figure S1-C). After particle extraction from motion corrected and Contrast Transfer Function (CTF) fitted images subsequent series of iterative 2D classifications facilitated the categorization of various protein classes (Figures S1-C). Collectively, the implementation of this method enabled the determination of cryo-EM structures for four different enzymes, achieving resolutions spanning from 2.7 to 4.2 Å. Considering the proteomic analysis, these enzymes were identified as dihydropyrimidinase-related protein 2 (DPYSL2), glutamine synthetase (GS), filamentous β-actin (F-actin) and V-ATPase. Cryo-EM image processing pipeline which contributed to each of these maps is shown in (Figure S2, S5, S7 and S9). In order to determine whether these four reconstructed proteins exhibit distinct or collaborative roles, we have used the STRING database. Our analysis revealed that all four proteins are involved in various interactions within this network (Figure S11).

### 2) DPYSL-2

The family of cytosolic phosphoproteins known as Collapsin Response Mediators (CRMPs), also referred to as DPYSL proteins, consists of five members and is highly expressed during nervous system development^16^. Here, we identified that the size exclusion chromatogram (SEC) fraction corresponding to mid-size proteins (440-150kDa) in our whole brain sample contains the enzyme DPYSL2, which was further confirmed by being the protein with the most PSMs (Figure S1-D).

Given that the corresponding particles of this protein were among the most populated 2D classes identified from the blob-picked particles, we were able to perform an ab-initio reconstruction. This was followed by heterogeneous refinement, which further cleaned up the particle stack, allowing us to achieve a 3D reconstruction using non-uniform refinement. We performed reconstruction both with and without enforcing symmetry. Consequently, we successfully determined its cryo-EM structure at a resolution of 2.7 Å with symmetry applied (D2 map) and 2.9 Å without symmetry (C1 map) (Figure S2).

Similar to the crystal structure of human CRMP2^21^, our reconstruction shows a tetrameric complex in which the monomer structure primarily adopts a triose-phosphate isomerase (TIM) barrel fold. Additionally, a distinct small β-sheet domain is formed by the N-terminal residues and segments near the C-terminus of the folded CRMP-2 (Figure 1-A). Each subunit of the tetramer consists of 572 amino acids, yet there exists a strong likelihood that approximately the final 100 C-terminal residues extend as unstructured chains from the central core of the tetramer^13,21^.

**Figure 1.**
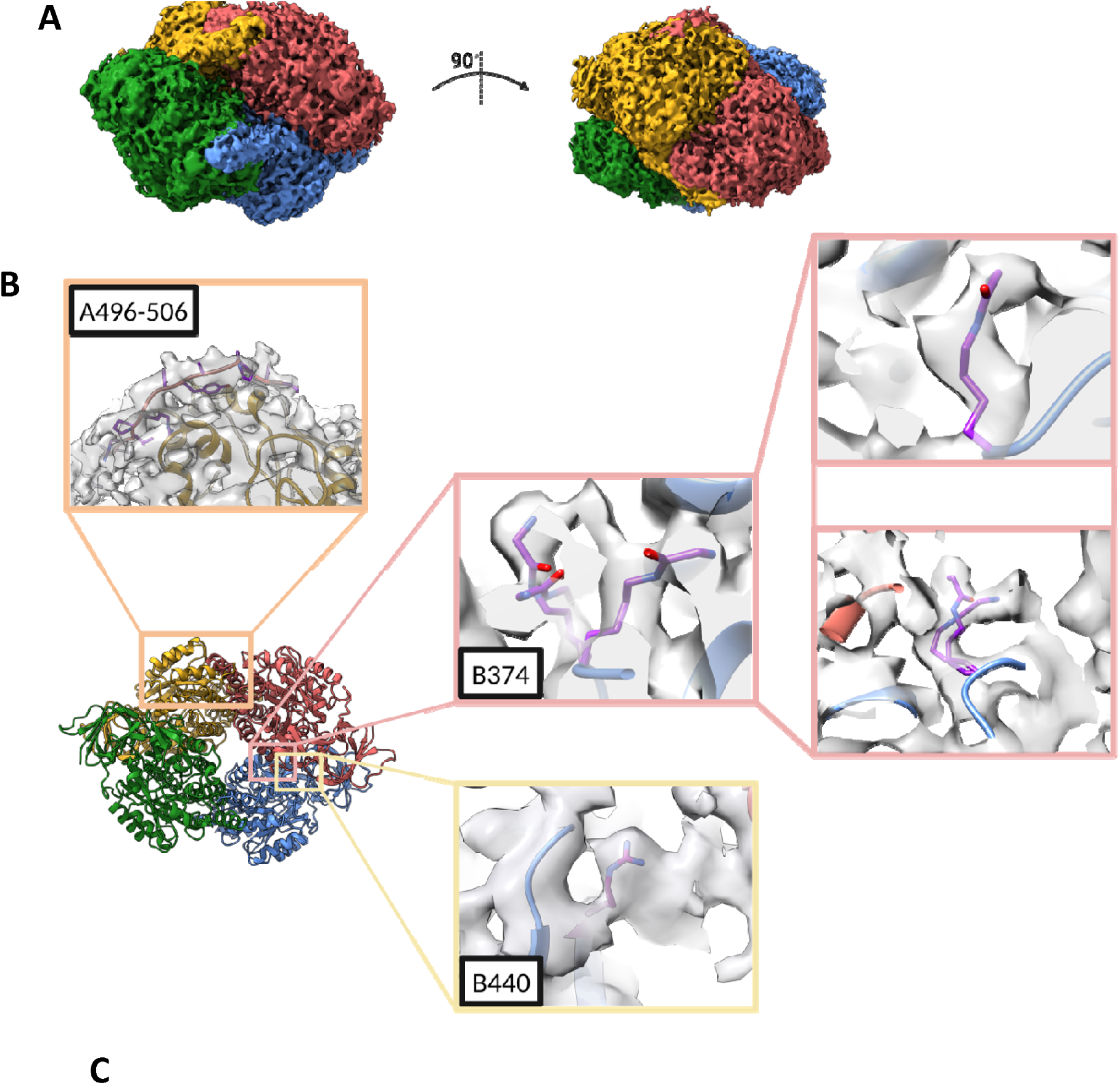
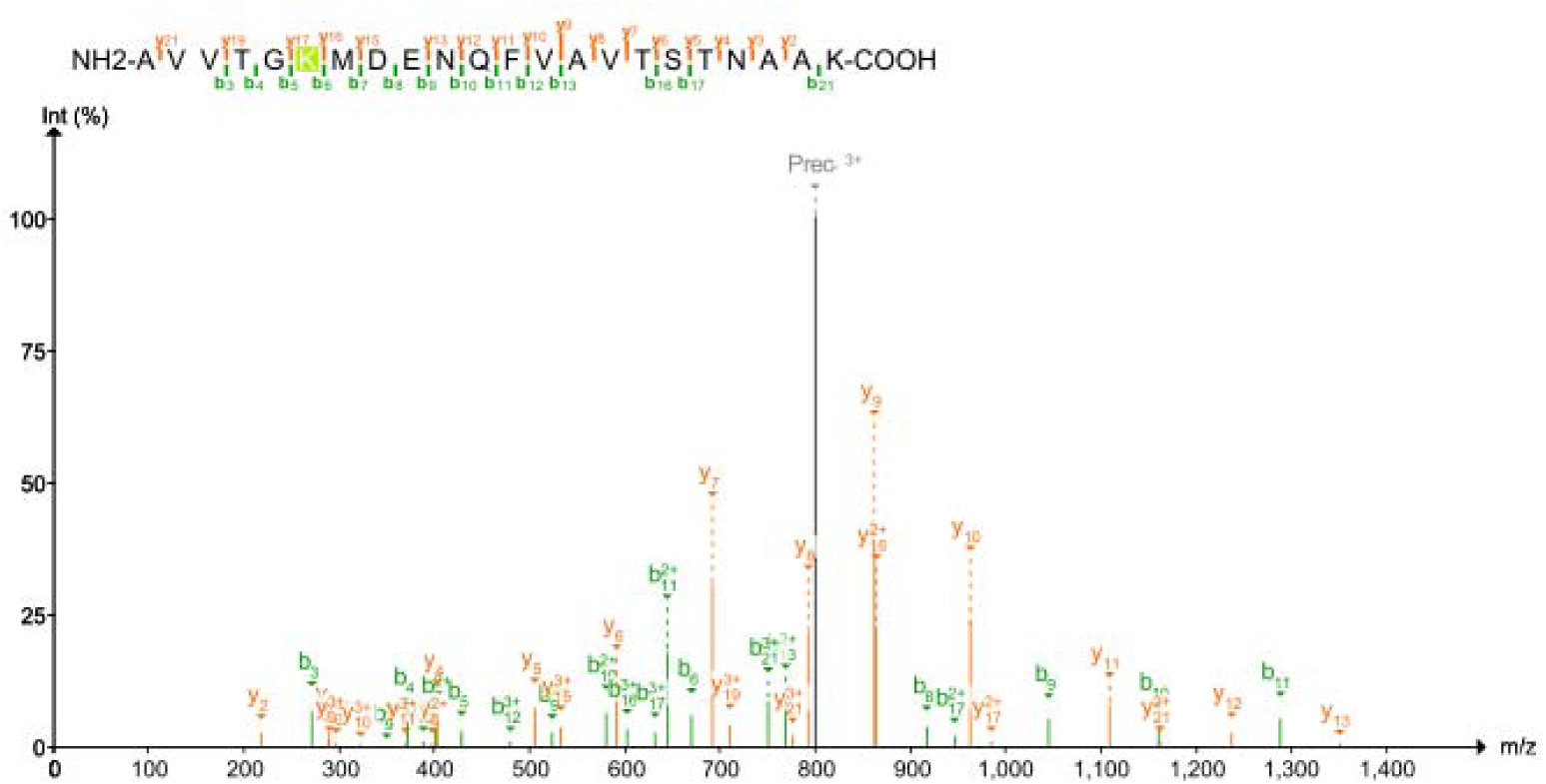
Structure of DPYSL2. **(A)** Cryo-EM map at 2.7 Å resolution of DPYSL2. Density corresponding to each protomer is colored differently. **(B)** Atomic model of DPYSL2 showing the tetrameric structure of the protein with D2 symmetry. Extra density labelled on chain A and chain B including C-terminus extra density and model up to residue 506 (orange), B374 with lysine conformations and SUMOylation related Glycine (pink), and extra density at Ubc9 interaction site (yellow). **(C)** MS spectra of the detected peptide relative to SUMO1 C-terminus Glycine-Glycine density on B374.

Previously, Zheng et al. successfully crystallized CRMP2, with an expression construct that encoded up to residue 526 of the protein. However, in their crystal structure, the electron density is predominantly visible up to residue 507 in only one of the four chains, indicating flexibility in this region^22^. Our findings align with this observation; as in our asymmetrically refined cryo-EM density map, we effectively modelled one of the chains up to residue 506 (Figure 1-B).

As of now, it is confirmed that DPYSL proteins transmit signals for multiple intracellular and extracellular pathways and are pivotal in various cellular functions such as cell migration, neurite extension, axonal guidance, dendritic spine development, and synaptic plasticity-all regulated by their phosphorylation status^16,23^. The latest findings indicate that CRMPs play crucial roles in various human pathologies, particularly neuropsychiatric, neurological, and neurodegenerative disorders^24^. It has previously been shown that phosphorylation of CRMP2 by Fyn kinase at tyrosine 32 (Y32) inhibits its SUMOylation at lysine 374 (K374). In contrast, when cyclin-dependent kinase 5 (Cdk5) phosphorylates CRMP2 at serine 522 (S522), it promotes SUMOylation at K374^25^. In our mass spectrometry data, we provide evidence of possible phosphorylation at S522 (Table S1) and high confidence of phosphorylation localized to S507, T509, S517, and S518. In our C1 cryo-EM density map of DPYSL2, we observe several unmodelled extra densities that can be related to some post-translational modifications (Figure S3) as well as three extra densities corresponding to different conformational states of SUMOylated K374 (Figure 1-B). We provide orthogonal evidence of SUMO1 C-terminus tail via a 114.0923 mass shift – corresponding to addition of GlyGly – localized to K374 in our mass spectrometry data (Figure 1-C and S4). It should be noted that, in AD, the accumulation of Aβ leads to an elevation in CRMP2 phosphorylation by Cdk5 at S522, which results in the depletion of CRMP2 from the microtubule network, disrupting various fundamental neuronal functions^26^.

Earlier studies affirm the potential of the CRMP2 SUMOylation site as a promising target for drug development and a direct approach is required to target the SUMOylation pocket surrounding residue K374 in CRMP2. This pocket is conserved among other members of the CRMP family across various species^27^. Nevertheless, it’s important to note that the key residue arginine 440 (R440), essential for anchoring CRMP2 to Ubc9, is present only in CRMP isoforms 2, 3, and 4. We observed cryo-EM densities located in the surrounding environment of this residue (Figure 1-B).

### 3) Actin

The actin cytoskeleton is essential for myriad cellular processes including transport, endo-and exocytosis, and cell division. Actin exists in two main forms: globular (G-actin) and filamentous (F-actin), which undergo continuous polymerization and depolymerization. F-actin predominates in biological activities, forming a network known as the actin cytoskeleton ^28,29^. Actin filament dynamics play a crucial role in shaping dendrites and synapses, contributing to their formation and remodeling^19^. Furthermore, researchers have noted the involvement of actin filament-related processes in enrichment analyses of both AD and bipolar disorder^30,31^

In this study, we present a structure of endogenous actin filaments extracted from AD rat brain cytosolic lysate with a resolution of 3.4 Å (Figure 2 and S5). To isolate the actin filaments, we subjected the filtered cytosolic brain lysate to size-exclusion chromatography. We then collected the fraction corresponding to 600 kDa< for freezing the grids containing the actin filaments. After performing the data acquisition, we manually picked some of the actin filament particles and then subjected their relative 2D classes to a Topaz train model^32^ in order to enhance the number of particles extracted from the data set (Figure S5). Followed by ab-initio reconstruction we conducted a helical reconstruction approach with a refined helical symmetry considering –166.77° rotation and 27.5 Å rise per subunit^29^ (Figure S5).

**Figure 2.**
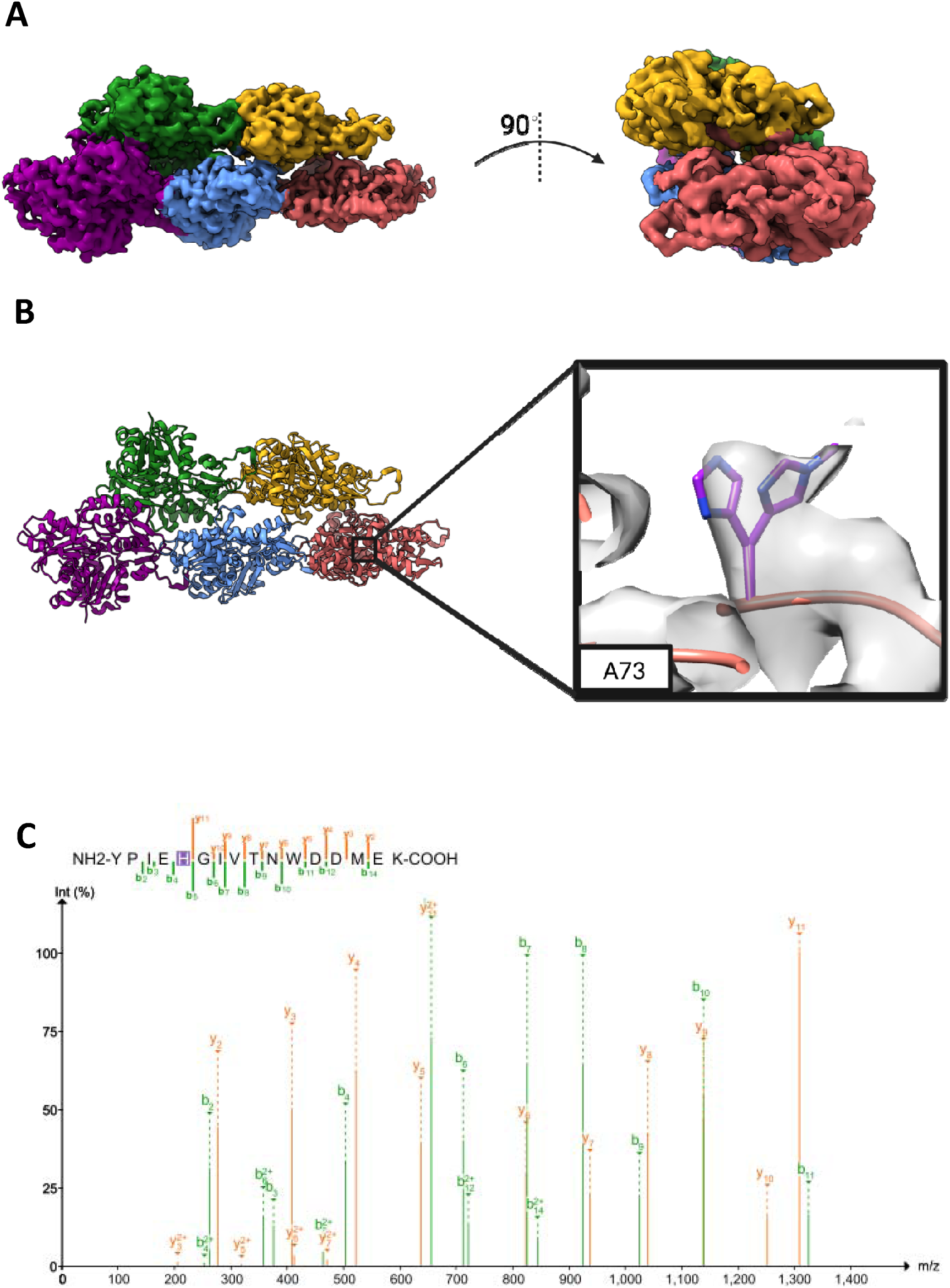
Structure of ACTIN filament. **(A)** Cryo-EM map at 3.4 Å resolution. Density corresponding to each actin protomer is colored differently. **(B)** Atomic model of Actin filament showing the helical structure of the protein. Zoomed view of the extra density located at H73 related to its methylation. **(C)** MS spectra of the detected peptide relative to Histidine 73 methylation.

The overall actin filament structure comprises four subunits (SD1-SD4) (Figure 2). In line with typical actin filaments, the brain filament derives stability from the interplay of actin subunits within the same strand (intrastrand) as well as those across opposing strands (interstrand). Examining the arrangement of F-actin is crucial in comprehending how it works. Since 2010, cryo-EM structure analyses of F-actin have been carried out^28,29,33^; however, to the best of our knowledge, this structure of F-actin is the first one captured in filament state without performing the polymerization in preparation.

Moreover, histidine methylation, a modification previously identified on a select few proteins, has been observed on actin, specifically at position H73 (or equivalent), across a diverse range of organisms, including worms, plants, and humans^34,35^. This modification is catalyzed by the methyltransferase enzyme SETD3, which is conserved among these organisms. Therefore, the presence of actin-His73me signifies an evolutionarily conserved modification found across a wide spectrum of multicellular eukaryotes^36^. The role of histidine methylation in modulating actin polymerization and smooth muscle contraction in mammals has been clarified, indicating its wide-ranging impact on the regulation of mammalian proteomes^36^. Here, in our refined 3D map, we identified an additional density corresponding to H73, indicating the presence of methyl group at this residue which we further confirmed with our MS analysis (Figure 2B, 2C and S6).

Mutagenesis investigations of Actin have shown that the triple mutant Q137A/D154A/H161A-actin lacks measurable ATPase activity^37^. Structural analyses of rabbit skeletal alpha actin have revealed the ATP-bound state at the specified residues in cryo-EM structures^38^. In our model of cytoplasmic F-Actin we can observe the densities relative to this site; however, no density for substrate could get resolved (Figure S7).

### 4) Glutamine Synthetase

In the brain, Glutamine synthetase (GS), which is specifically expressed in astrocytes, forms glutamine by an ATP-dependent amination of glutamate. Various pathological factors associated with Alzheimer’s disease can directly diminish the activity of glutamine synthetase, including Aβ deposition, chronic inflammation, hypoxia, ischemia/reperfusion, and oxidative stress^39,40^.

Although it has been shown that 3xTg-AD mice exhibit a decrease in GS astrocyte expression, glutamine synthetase was amongst the proteins with the most PSMs which contributed to 12098 particles in our cryo-EM density and by applying the BaR method after inducing the D5 symmetry we were able to reconstruct its 3D volume to 3.1 Å (Figure 3 and S8).The overall structure of the GS, composed of two doughnut-shaped pentamers in which 10 identical subunits are stacking into two layers (D5 symmetry) (Figure S8). These two layers are connected by flexible loops which consist of 11 residues each (103-113).

**Figure 3.**
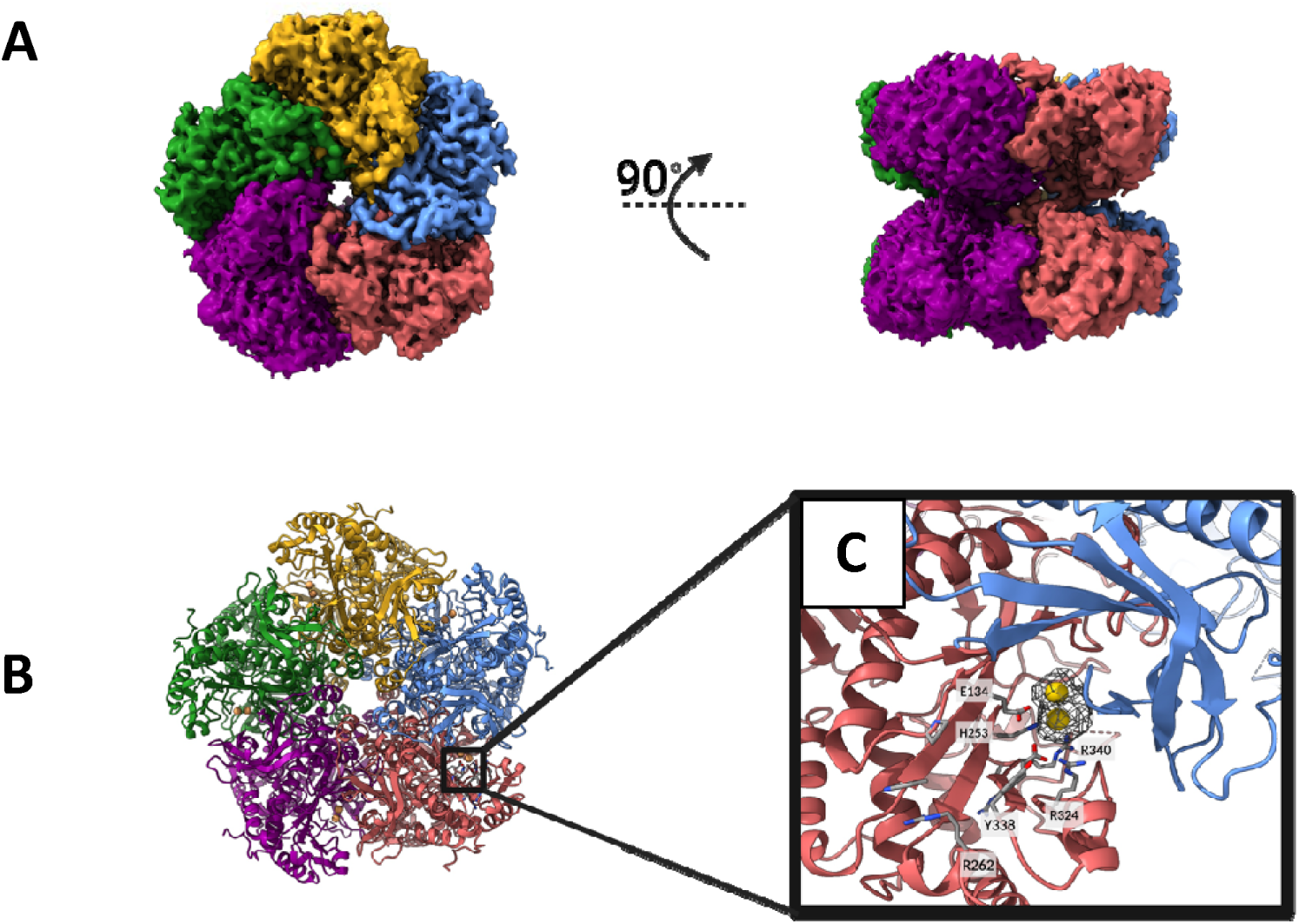
Structure of Glutamine Synthetase. **(A)** Cryo-EM map at 3.1 Å resolution. Density corresponding to each protomer is colored differently. **(B)** Atomic model of Glutamine Synthetase showing the 10 mer structure with D5 symmetry.**(C)** Zoomed view of Mn^2+^ binding site with surrounding charged residues.

The N-terminal section which is relatively smaller than the C-terminal consists of four alpha-helices and five beta strands which is responsible for securing the inter-subunit interactions within the layer of pentamer. The C-terminus is composed of seven alpha helices and seven beta-strands in which residues 149-157 are responsible for anchoring the pentameric stack layers into each other. The sequence alignment with Bovine (Bos taurus) glutamine synthetase revealed a 93.3% identity with the rat counterpart (Figure S9-A). When superimposed with the structure from Bovine^13^,(PDB ID:7U5N), our model exhibited a minimal RMSD of 0.03 Å (for 337 Cα atoms). Notably, disparities emerge in the C-terminus between our structure and the bovine counterpart. Illustrated in Figure S9-B, small alpha helices are present within the connecting C-terminus loops. Despite complete conservation of the sequence (residues 291-299) at this position between bovine and rat, these helices are absent in the bovine model.

Within a GS decamer, there are ten active sites distributed at the juncture where two adjacent subunits connect within the pentameric ring. The N-terminal β-sheet domain of one subunit interacts with the highly curved β-sheet found in the C-terminal catalytic domain of the neighboring subunit. This interaction gives rise to the creation of active pockets^41^. The vicinity of this site also contains multiple positively charged arginine residues, such as R262, R324, R340, and R341, which play a role in substrate binding^13^. In our cryo-EM density map, we identified two Mn^2+^ ions, coordinated by residues E134, H253, and E338 through polar interactions (Figure 3-B).

### 5) V-ATPase

Vesicular- or vacuolar-type adenosine triphosphatases (V-ATPases) are proton pumps driven by ATP hydrolysis that are essential for acidifying endosomes, lysosomes, and the trans-Golgi network, as well as for acid secretion by osteoclasts, kidney intercalated cells, and some tumor cells^42^. In brain cells, the loading of neurotransmitters into synaptic vesicles relies on the energy provided by proton-pumping V-ATPases^43,44^.

The V-ATPase enzyme consists of soluble V_1_ and membrane-embedded V_O_ regions. ATP hydrolysis by A and B subunits in V_1_ drives rotor rotation, connected to V_O_ through stalks. This rotation facilitates proton translocation across the membrane via specific channels, linking ATP hydrolysis to proton transport. Cryo-electron microscopy has revealed three distinct conformations of the enzyme, termed ’State 1’, ’State 2’, and ’State 3’, even in the absence of ATP^45,46^.

Proper function of the V-ATPase, vital for its rapid ATP hydrolysis and proton transport, necessitates tight regulation. For instance, although V-ATPase activity is crucial for loading secretory vesicles, its activation at the plasma membrane during exocytosis is undesired. To maintain control over V-ATPase activity, the regulated separation and subsequent reassembly of its V_1_ and V_O_ regions are essential mechanisms^45,47^. Throughout the process of ATP hydrolysis and proton pumping, the intact V-ATPase undergoes sequential cycling between three distinct rotational states: States 1, 2, and 3^46^.

When glucose levels decrease, the V_1_ and V_O_ regions of the V-ATPase enzyme disengage or separate^45^. During this separation it has been shown that V complex is adopting State 3 of rotation while V_1_ complex is adopting State 2. It remains unknown whether the separation of the V-ATPase occurs during State 2 or State 3 of its rotational cycle. In State 2, the V_1_ structure indicates an inability to shift to State 3 without the dissociation of subunit C. Once subunit C separates, V_1_ΔC gains the capability to explore all three rotational states^46^.

To obtain highly purified samples for single-particle image analysis on cryo-EM datasets, researchers are leveraging the previously described rotation states of V-ATPases bounded with Sid-K^43^. Here in our cryo-EM dataset of cytosolic brain proteins we were able to identify V_1_ V-ATPase particles presented in our sample. After performing the BaR method aiming to reconstruct the V-ATPase 3D structure, we faced preferred orientation of this endogenous V_1_ V-ATPase in our sample which contributed to missing views for the side panel and anisotropic 3D map. However, the quality of this map was sufficient that enabled us to dock different state V_1_ V-ATPase models. Performing real space refinement for our V-ATPase map against reported state2 (PDB ID: 7TMP) at 4.2 Å suggested that the V_1_ reconstruction we could capture was arrested in State 2 (Figure 4 and S10) which was previously shown to be the state in which V_1_ region of V-ATPase gets dissociated from its membrane embedded region of V_O_.

**Figure 4.**
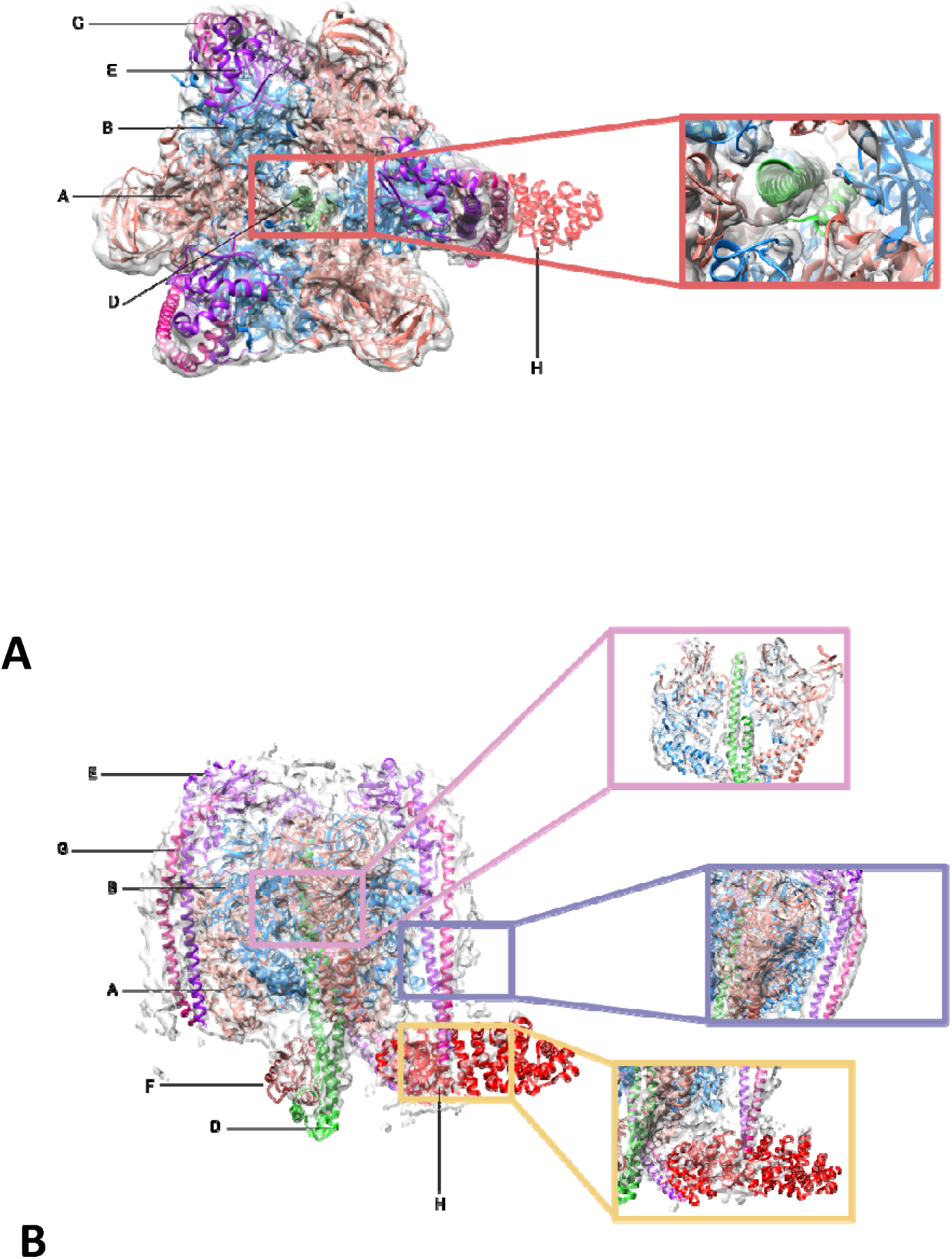
Structure of V_1_ V-ATPase. **(A)** Top view Cryo-EM map of fully assembled V_1_ at 4.24 Å resolution. 7TMP PDB atomic model docked into the density map, and it is suggesting State 2 of V_1_ rotation considering the subunit D. **(B)** Side view Cryo-EM map of fully assembled V_1_ at 4.03 Å resolution. 7TMP PDB atomic model docked into the density map, and it is suggesting State 2 of V_1_ rotation considering the subunits G, E, H, and D.

Moreover, in our particle stacks obtained from the Topaz-trained model for picking V-ATPase particles, we noticed some particles representative of half-assembled or truncated V_1_ regions. We refined these particles to perform docking of V_1_ V-ATPase models; however, it seems that this reconstruction is an average of all 3 rotational states (Figure S10-B).

## Discussion

Given the complex and multifactorial nature of AD, there is an imperative for the identification and validation of novel therapeutic targets to address the current gaps in treatment efficacy and disease modification. Capturing potential target protein structures in their native state is paramount for understanding the intricate molecular mechanisms underlying AD pathology, as it provides invaluable insights into protein-protein interactions, post-translational modifications, conformational changes, and functional dynamics crucial for target identification and drug development. The BaR methodology offers a means to uncover structural details of diverse protein constituents within a raw biological sample, achieving near-atomic resolution^12^. The BaR method has previously been utilized to investigate protein complexes extracted from human brain microsomes^15^. However, the utilization of brain microsomes may present limitations in offering a native protein environment, intricate protein-protein interactions, a wider array of proteins, and a more accurate representation of cellular processes.

In this study, we employed the BaR method in conjunction with mass spectrometry analysis to elucidate the structural proteome of a rat brain model of AD. This approach facilitated the 3D reconstruction of four protein complexes in their native states, encompassing substrate-bound conformations, post-translationally modified residues, and arrested states. Interestingly, all these protein complexes are linked to the pathology of neurodegenerative disorders including AD, PD and HD.

It has been shown that, Phosphorylation of CRMP2 by Cdk5 at S522^48^ primes it for subsequent phosphorylation by GSK3β at T509/514/518^49^ and this form of CRMP2 has been implicated in various disorders such as bipolar disorder^50^, Alzheimer’s disease^24^, and certain cancers. For example, In AD, CRMP2 is phosphorylated by enzymes GSK3β and CDK5, like tau protein, leading to neurofibrillary tangle formation. Phosphorylated CRMP2 is found in NFTs along with the synaptic structure-regulating SRA1/WAVE1 complex, potentially contributing to neural and synaptic structure deficits seen in AD^51^. Additionally, Cdk5-phosphorylated CRMP2 can undergo SUMOylation by Ubc9 at K374, a process that can be hindered by CRMP2 phosphorylation by the src kinase Fyn at Y32^27^. In conditions of chronic pain, levels of phosphorylated and SUMOylated CRMP2 are elevated, allowing it to modulate the activity of nociceptive ion channels such as CaV2.2 and NaV1.7. Furthermore, the association of CRMP2 with CaV2.2 contributes to behaviors associated with cocaine relapse. Interestingly, in our mass spectrometry and cryo-EM analysis we could find the evidence of phosphorylated T509 and S517/518, S522 as well as K374 SUMOylation^52^.

Moreover, pull-down investigations have yielded a rich assortment of proteins that either directly interact with CRMP2 or assemble into multimeric complexes alongside CRMP2. Interestingly, actin in its all native forms (monomeric/ filamentous) is amongst the highest interaction partners for CRMP2^51,53^. In our data, we also have the validation of actin abundance in our sample through mass spectrometry analysis and cryo-EM imaging.

The function of the actin cytoskeleton in dendritic spines stands out as a key biological pathway essential for both synaptic plasticity and synaptic dysfunction during the initial phases of AD^54^; Hence, obtaining the natural structure of an actin filament from an AD model could prove instrumental in gaining deeper insights into its role in neurogenesis.

Additionally, we observed extra density, likely indicating the presence of a methyl group on H73. Previous structural analyses, supported by biochemical experiments and enzyme activity assays, have shown that SETD3 exhibits high specificity in recognizing and methylating beta-actin. Furthermore, both SETD3 and beta-actin undergo significant conformational changes upon binding to each other^55^.

GS plays a role in converting glutamate to glutamine, mitigating the toxicity of glutamate at levels that could cause neuronal damage. A recent study found that individuals with probable AD or major depression had higher levels of glutamate and glutamine in their cerebrospinal fluid^56^. This can be further approving an earlier study which has been shown that glutamine synthetase activity is significantly decreased in AD which supports the hypothesis of an accumulation of oxidized (inactivated) glutamine synthetase^57^.

In V-ATPase, Reports indicated that a neuronal degradation mechanism dependent on V0a1 may be causally associated with AD pathology^58^. In mammalian V-ATPase structure, there are two transmembrane alpha helices that are located within the c ring in V_O_ region^43^. It has been shown that mutations in these two helices can cause cognitive impairment, X-linked parkinsonism and epilepsy^59,60^. Utilizing the BaR method, we can focus on extracting multi-subunit complexes such as V-ATPases from various disease sources.

In this study, despite encountering obstacles, we were unable to resolve the atomic model of the V_1_ region within this protein complex. Nevertheless, our findings remain significant as through our 3D map reconstruction, it appears that a majority of the particles have been arrested in a State 2 of rotation while we could not observe the density relative to subunit C. This might be due to loss of this subunit during sample preparation or lacking enough sideview that prevented us to resolve the density for it in our 3D map. This observation holds promise for elucidating structural alterations in the endogenously untagged purification of this protein, offering valuable insights for further investigation.

Considering that all the proteins discussed in this study are linked to the pathological state of the Alzheimer’s model utilized, it sheds light on the potential and feasibility of employing the BaR methodology to surmount the limitations of cryo-EM in heterogeneous and impure samples. Such an approach holds promise for delving deeper into the intricate structural nuances of proteins present in cancer tissues or pathological blood plasma, facilitating further investigation into disease mechanisms and potential therapeutic targets.

## Materials and Methods

### 1) Animal model

TgF344-AD rats are outbred on a Fischer strain and housed on a 12-hour light:12-hour dark schedule with ad libitum access to food and water. Ethical approval of all experimental procedures was granted by The Animal Care Committee of the Sunnybrook Health Sciences Center, which adheres to the Policies and Guidelines of the Canadian Council on Animal Care, the Animals for Research Act of the Provincial Statute of Ontario, the Federal Health of Animals Act, and is compliant with ARRIVE guidelines.

### 2) Protein purification

A 22-month-old female TgF344-AD rat brain was deeply anesthetized with 5% isoflurane and 1% O_2_ and subsequently transcardially perfused with cold phosphate-buffered saline (PBS, Wisent) for 5 minutes. Following perfusion, the brains were promptly removed and stored in 50 mL of cold PBS on ice.

The tissue was then homogenized using a mechanical homogenizer in cold lysis buffer (comprising 20 mM HEPES, 0.5 mM PMSF, 1mM Benzamidine, 5mM aminocaproic acid, 10mM NaF, 50 mM β-Glycerophosphate at pH 7.4). After homogenization, the lysate was extracted and subjected to sequential centrifugation steps at 4000 xg for 10 minutes at 4°C, followed by 150,000 xg for 1 hour at 4°C.

The resulting cell lysate was filtered and then passed through a size exclusion column, specifically Superose 6 10/300 increase column, equilibrated with buffer containing 20 mM HEPES, 150 mM NaCl, 1 mM DTT, at pH 7.4. Fractions were collected based on their molecular weight, with particular attention to those within the range of 150-450 kDa, as well as earlier fractions containing higher molecular weight components such as actin filaments.

### 3) Cryo-EM sample preparation and data collection

3 μl of size separated protein mixture was applied onto in-house nanofabricated holy sputtered gold girds with a hole size of ∼2 μm and a 4 μm period. Grid freezing was done in Vitrobot Mark IV (FEI) with 3-second blotting at 4 °C, 90% humidity and using liquid ethane kept at liquid nitrogen temperature.

### 4) Data collection

A Titan Krios G3 microscope operated at 3001kV and equipped with Falcon 4i camera (TFS) was used to collect data in super-resolution mode at 92,000× magnification (physical pixel size of 1.030 Å/pix, 0.535 Å/pix super-resolution). Micrographs of the of each separated fractions were collected in two separate datasets (30 frames, total dose 40 or 41 e−/Å2) Automated data collection was performed by EPU (Table S2).

### 5) Image analysis

Patch motion correction and patch CTF estimation were used to align and averaged movie frames and correct for CTF in *cryoSPARCv4* ^61^ with last movie frame ignored. A modified BaR method was used for reconstruction of each protein from each sample^12^. First, automatic particle picking was done using blob picker job followed by particle extraction from curated movies. Series of iterative 2D classification were performed in order to clean up the particle stacks by manually selecting meaningful classes and repeating classification until no new clear classes were seen. Initial low-resolution maps for individual proteins were solved using iterative rounds of *abinitio*-3D reconstruction followed by Non-Uniform refinement in cryoSPARC. In cases of limited particle number we trained a topaz model^32^ using the relative 2D class templates followed by particle extraction through Topaz extract job and performing Non-Uniform refinement with applying symmetry. For reconstructing the Actin filament, after extracting the particles we performed a helix refinement in cryoSPARC with helical twist estimation of -167 degrees and helical rise estimation of 26.7Å.

### 6) Model building and refinement

Initial models of GS and DPYSL-2 were built via SWISS model performed by using the Uniprot sequence of each protein and aligned to the cryo-EM density maps in Chimera. For the filament 8OID used as the initial model to dock into the sharpen map. The ultimate models were manually refined using Coot and enhanced through the phenix.real_space_refine feature within phenix.

Ligands were added if there was clear density that matched previously described structural or biological data. Final structures were evaluated using MolProbity^62^. Statistics associated with data collection and refinement are compiled in Table S3. Final figures were generated using the ChimeraX suite^63^.

### 7) Proteomic sample preparation

Sample was brought to 5 mM Tris(2-carboxyethyl)phosphine and 100 mM ammonium bicarbonate and 20% acetonitrile was added to assist with digestion^64^. Samples were placed in an Eppendorf Thermomixer at 37 °C and allowed to reduce for 30 min at 800 rpm. For alkylation, sample was brought to 10mM iodoacetamide and incubated at 37 °C for 30 min at 800 rpm. Trypsin/Lys-C (Promega, V5072) was added to each sample at 1:20. Samples were allowed to digest for 16 h in a 37 °C at 800 rpm. Samples were cleaned using SP2^65^ with 4 µL of of prewashed magnetic particles (100 ug/uL of 1:1 SeraMag Hydrophilic : SeraMag Hydrophobic).

### 8) Mass spectrometry data acquisition

LC-MS/MS analysis was performed on an Orbitrap Eclipse Tribrid MS (ThermoFisher) coupled to Neo Vanquish Liquid Chromatography system (ThermoFisher). Peptides were loaded on pre-column (ThermoFisher, 174500) with 40 μL of mobile phase A (0.1% FA in HPLC grade water) at 30 μL/min and separated using a 50 cm EASY-Spray column (ES803, ThermoFisher) ramping mobile phase B (0.1% FA in HPLC grade acetonitrile) from 0% to 5% in 2 min, 5% to 30% in 103 min, 30% to 48% in 21 min interfaced online using an EASY-Spray™ source (ThermoFisher). The Orbitrap Eclipse Tribrid MS was operated in data dependent acquisition mode using a 3 s cycle at a full MS resolution of 240,000 with a full scan range of 350-1550 m/z with RF Lens at 60%, full MS AGC at 250%, and maximum inject time at 50 ms. Ions were accumulated using a 1.4 Th isolation window, 35 ms maximum injection time, with an AGC target of 200%. Ions for MS/MS were selected using monoisotopic peak detection, intensity threshold of 1,000, positive charge states of 2-5, 40 s dynamic exclusion, and then fragmented using HCD with 30% NCE. MS/MS scans were recorded in the Ion trap with Rapid Scan Rate over 200-1400 m/z.

### 9) Mass spectrometry raw data analysis

Raw files were analyzed using FragPipe (v.20.0) using MSFragger^66,67^ (v.3.8) and Philosopher v.5.0.0 ^68^.Data were searched against a human database including canonical plus isoforms (Uniprot, 43,392 sequences). Default settings, including ProteinProphet^69^ and Percolator^70^ for PSM validation, were used. For post-translational modification searches, each modification was searched separately, specified as variable modification – STY:79, H:14.01565, or K:114.0923, and localized with PTMProphet^71^. Images of spectra were generated using FragPipe-PDV^72^.

### 10) Interaction Observation

Using multiple protein function in STRING data base we ran a search for protein uniport gene names (#1 DPYSL2 #2 Glul #3 ACTB #4 Atp6v1b2). Interaction confidence was represented using variations in line thickness.

## Supplementary figures

**Figure S1.**
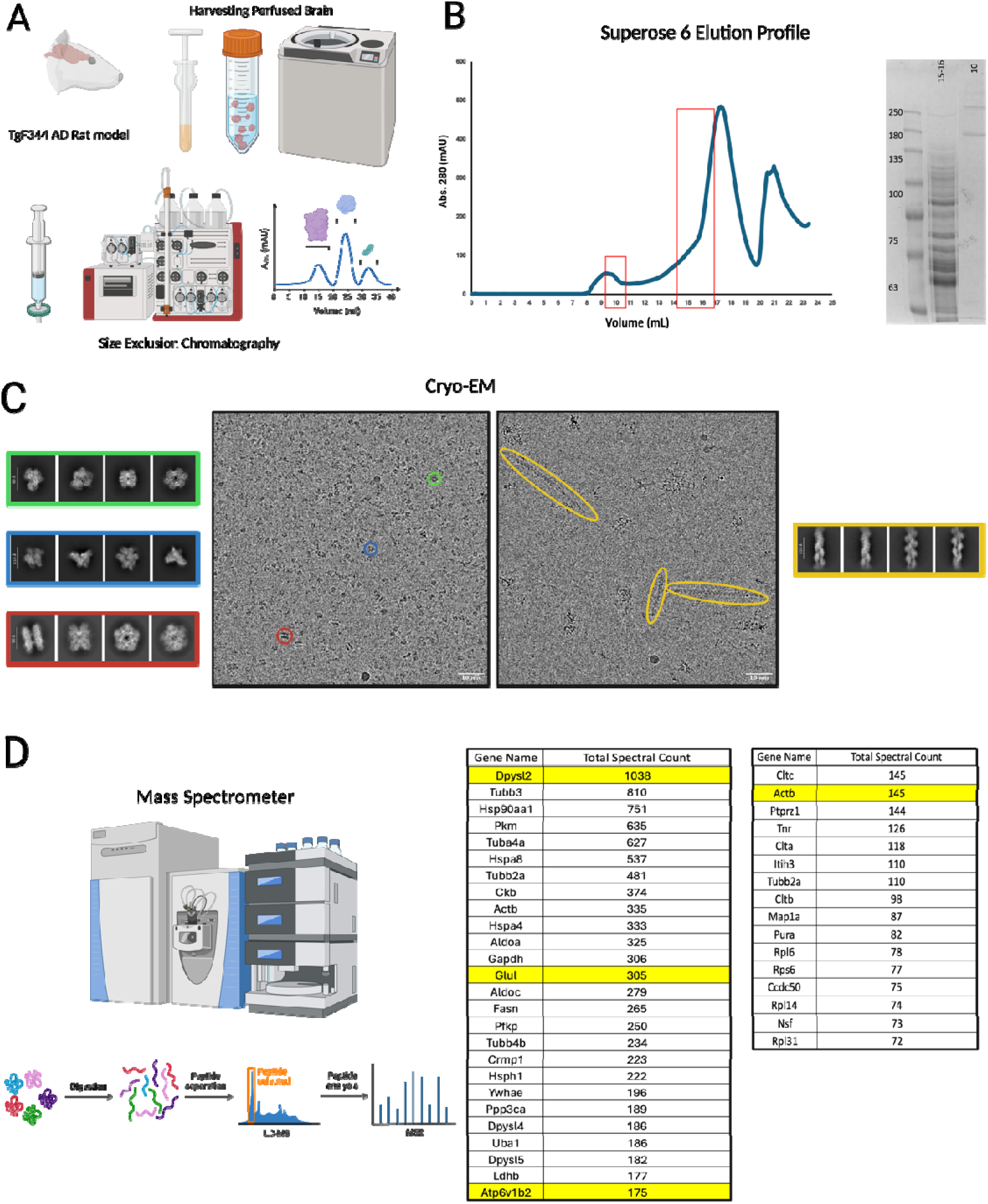
**(A)** Overview of the experimental method. **(B)** Size exclusion chromatogram (SEC) showing the elution profile and SDS page gel relative to SEC fractions. **(C)** Representative Cryo-EM micrographs showing the picked particles and their relative 2D-classes. Left and right panels correspond to fractions 15-16 and 10, respectively, shown in S1B. **(D)** Mass spectrometry overview and the tables of most populated proteins in each SEC fraction (Showing the relative structurally solved proteins in yellow highlight). Left and right tables correspond to fractions 15-16 and 10, respectively, shown in S1B.

**Figure S2.**
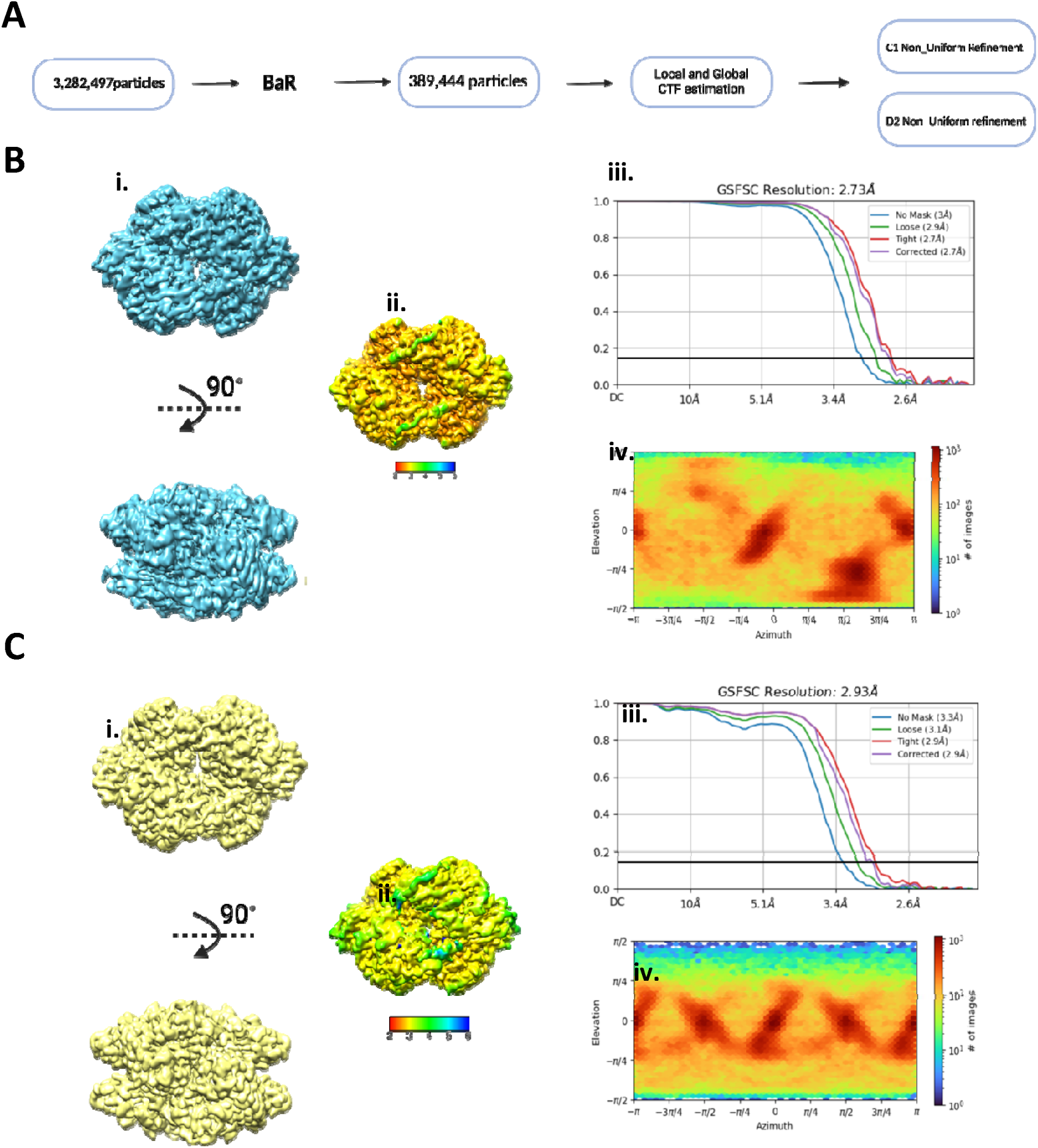
**(A)** Schematic overview of BaR analysis. **(B) i.** Reconstructed Cryo-EM density map of DPYSL2 with D2 symmetry. **ii.** Local resolution estimation map. **iii.** GS-FSC curve showing final resolution at 2.7Å. **iv.** Viewing direction distribution. **(C) i.** Reconstructed Cryo-EM density map of DPYSL2 with C1 symmetry. **ii.** Local resolution estimation map. **iii.** GS-FSC curve showing final resolution at 2.9Å. **iv.** Viewing direction distribution.

**Figure S3.**
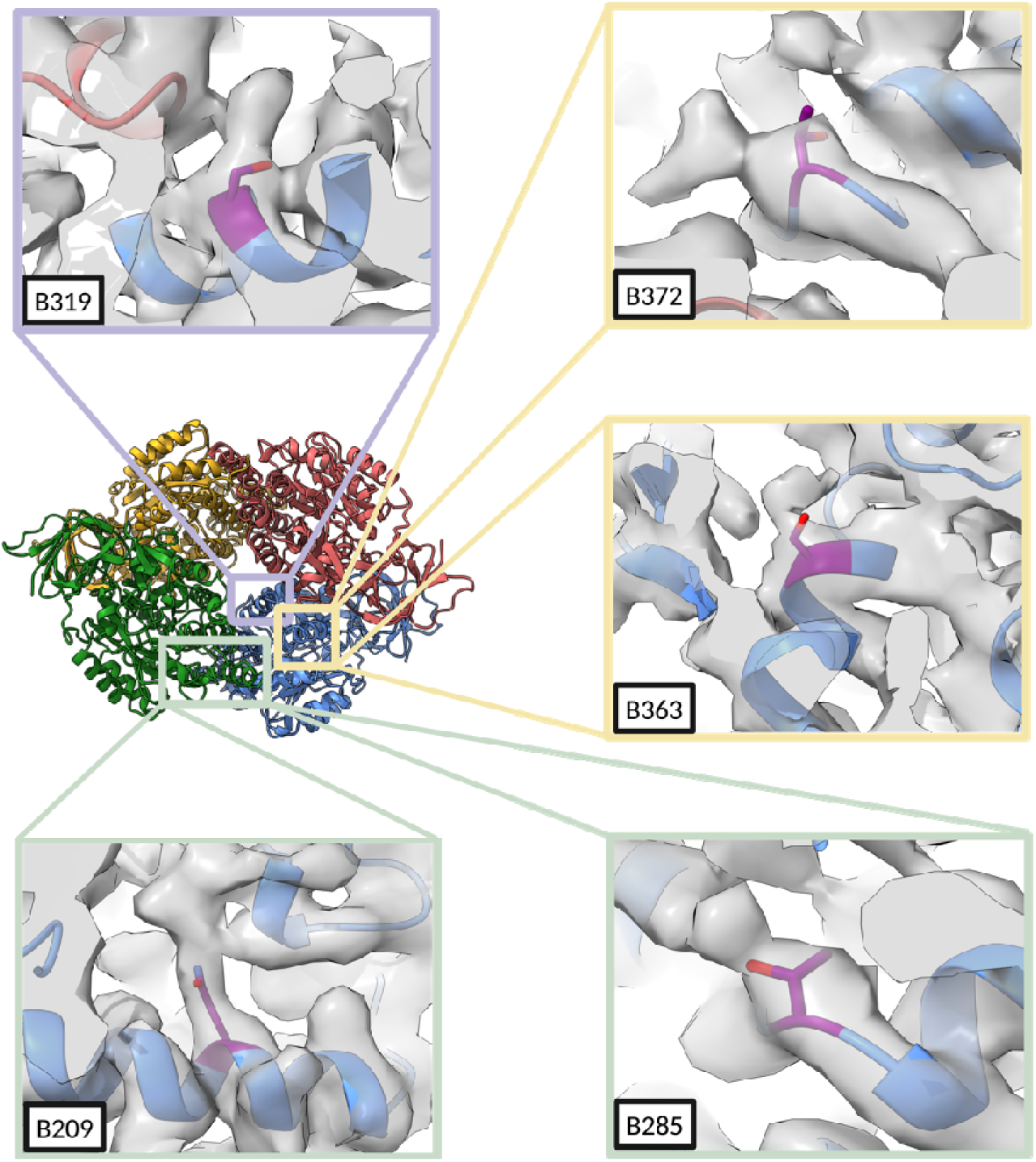
The extra densities on the DPYSL reconstructed map shown with atomic model that can be associated with potential phosphorylated residues.

**Figure S4.**
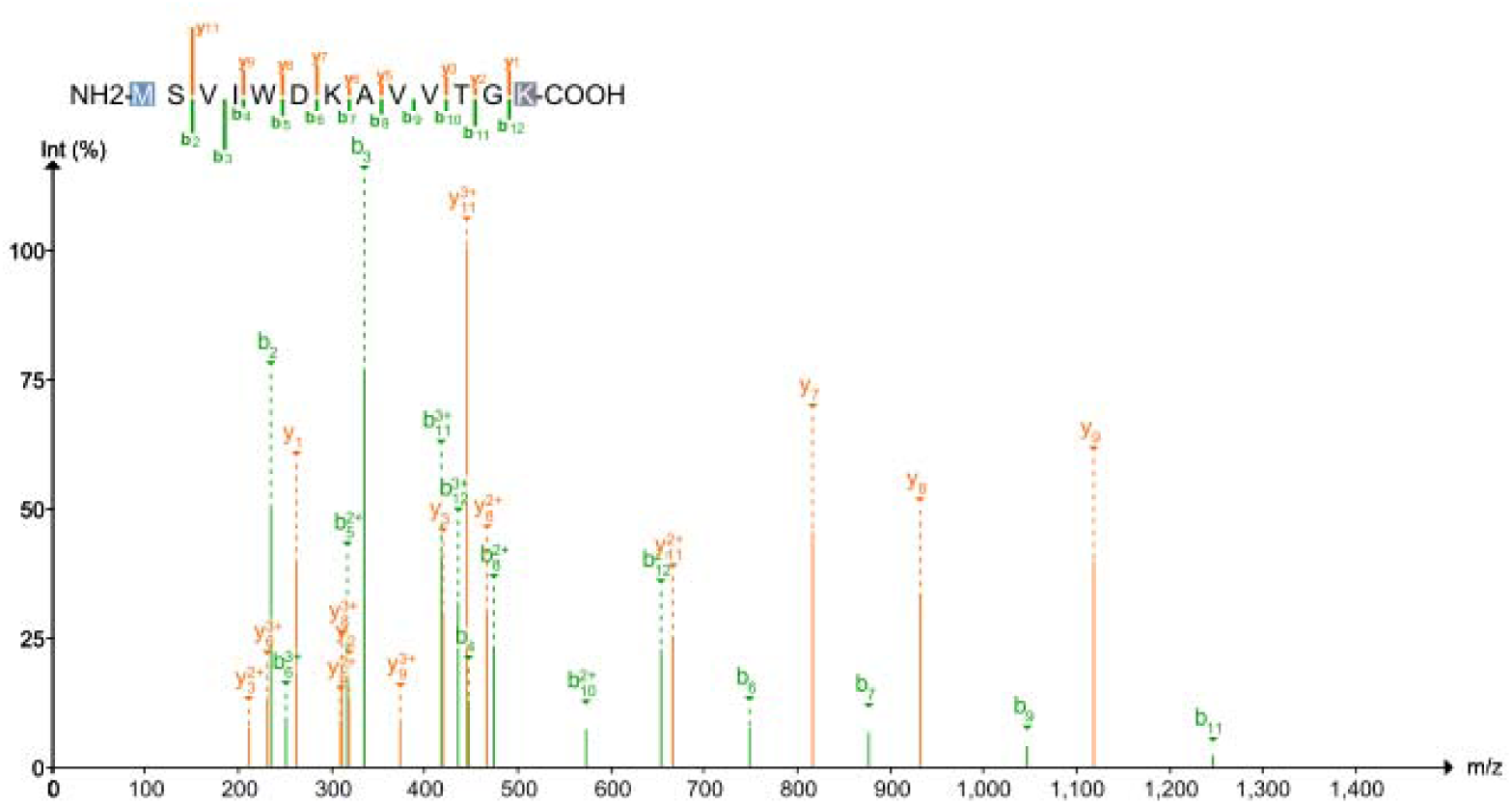
MS spectra of the detected peptide relative to SUMO1 C-terminus Glycine-Glycine density on B374 in DPYSL2.

**Figure S5.**
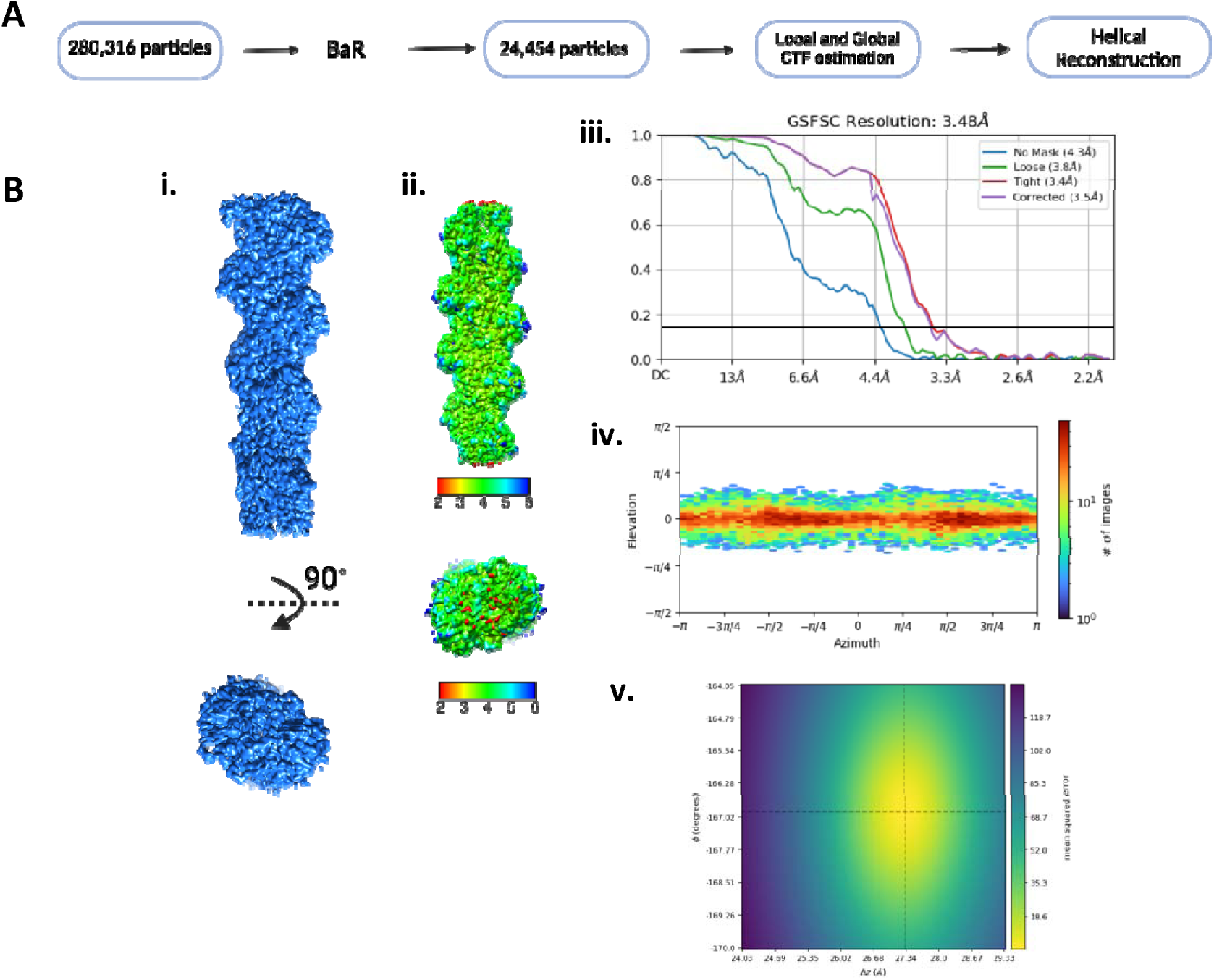
**(A)** Schematic overview of BaR analysis. **(B) i.** Reconstructed cryo-EM density map of Actin filament with helical symmetry. **ii.** Local resolution estimation map. **iii.** GS-FSC curve showing final resolution at 3.4Å. **iv.** Viewing direction distribution. **v.** Helical symmetry error surface.

**Figure S6.**
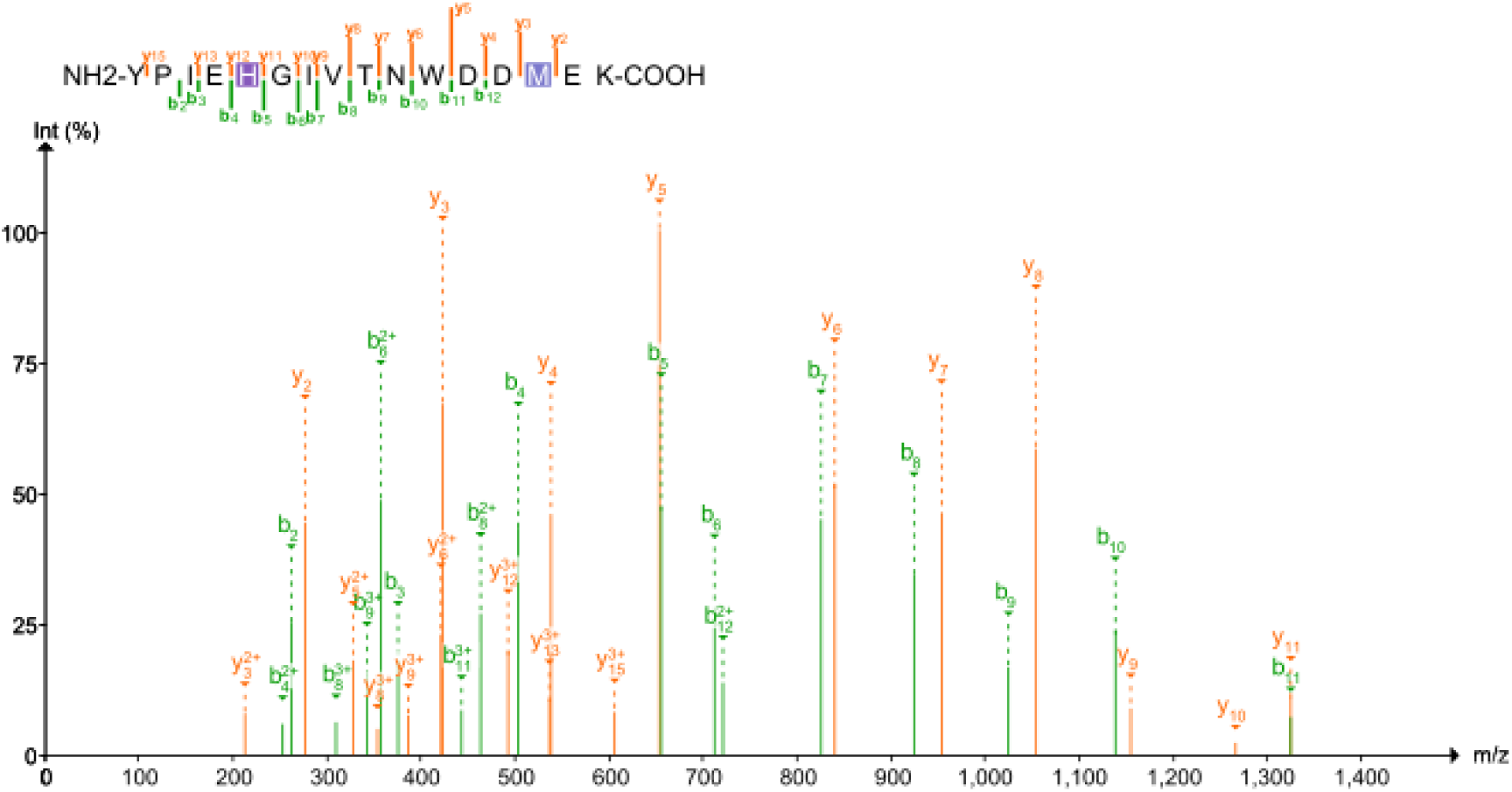
Mass spectrophotometry spectra of the detected peptide relative to methylated Histidine 73 in Actin.

**Figure S7.**
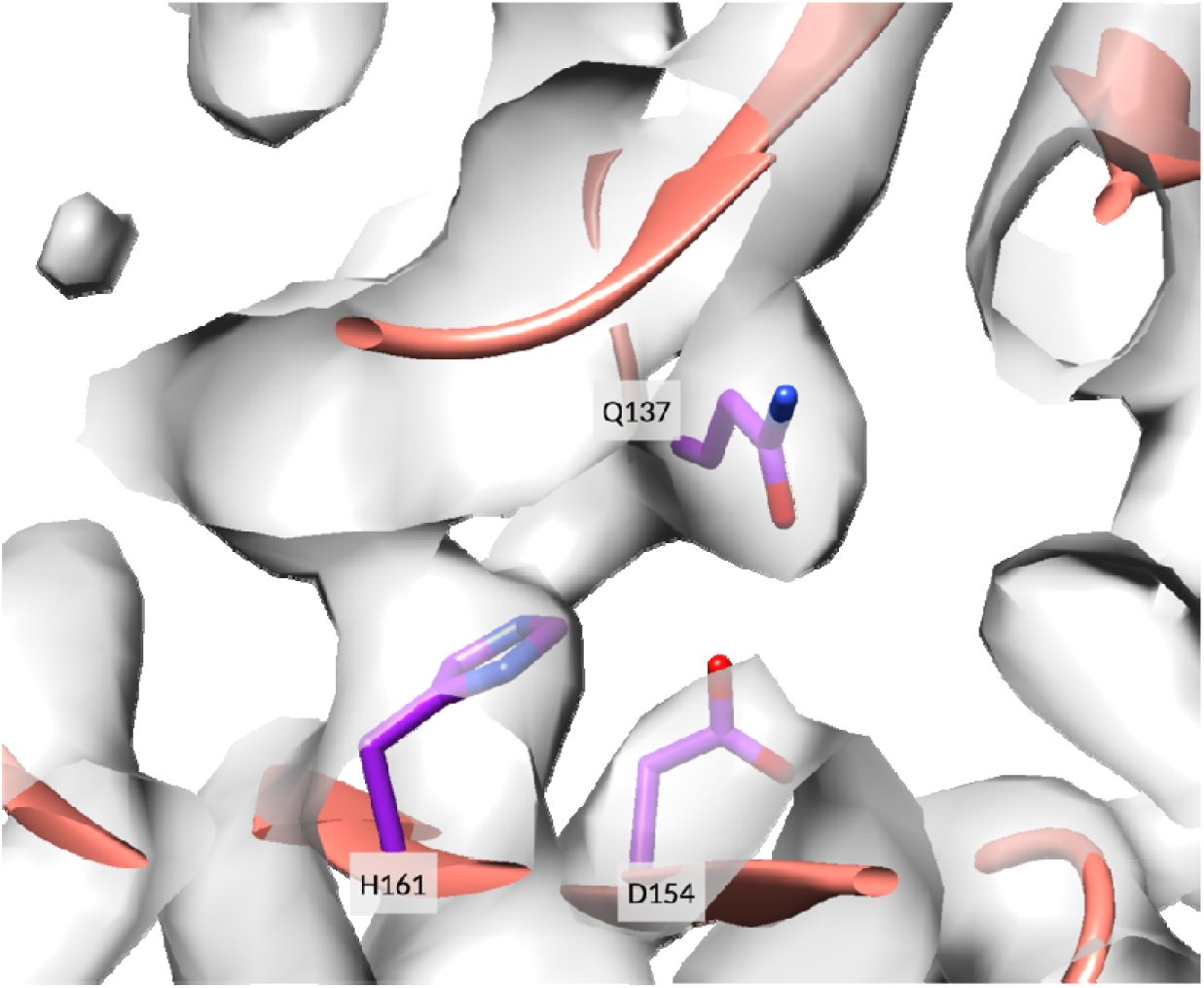
Actin substrate binding site.

**Figure S8.**
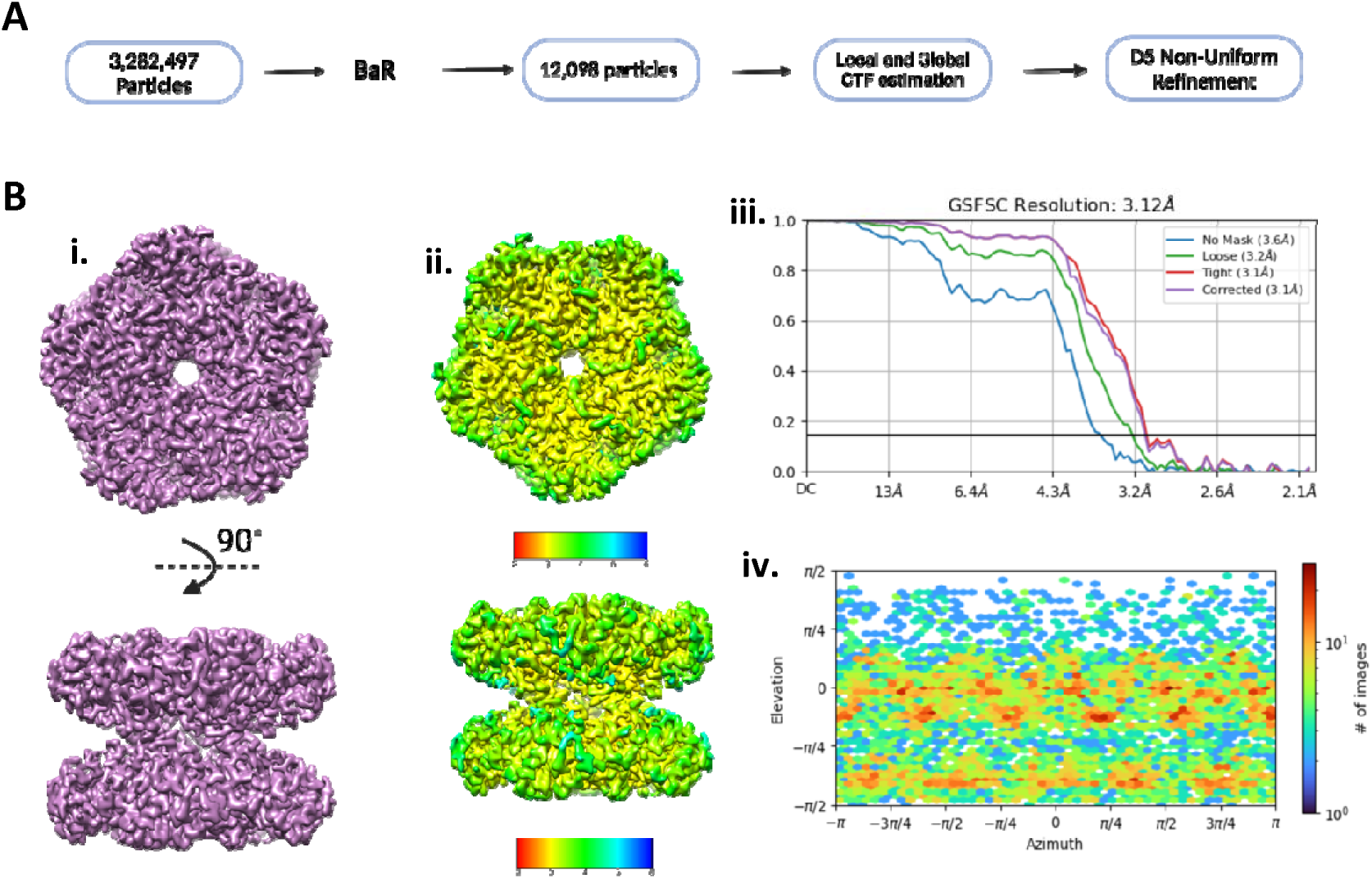
**(A)** Schematic overview of BaR analysis. **(B) i.** Reconstructed cryo-EM density map of Glutamine synthetase with D5 symmetry. **ii.** Local resolution estimation map. **iii.** GS-FSC curve showing final resolution at 3.1 Å. **iv.** Viewing direction distribution.

**Figure S9.**
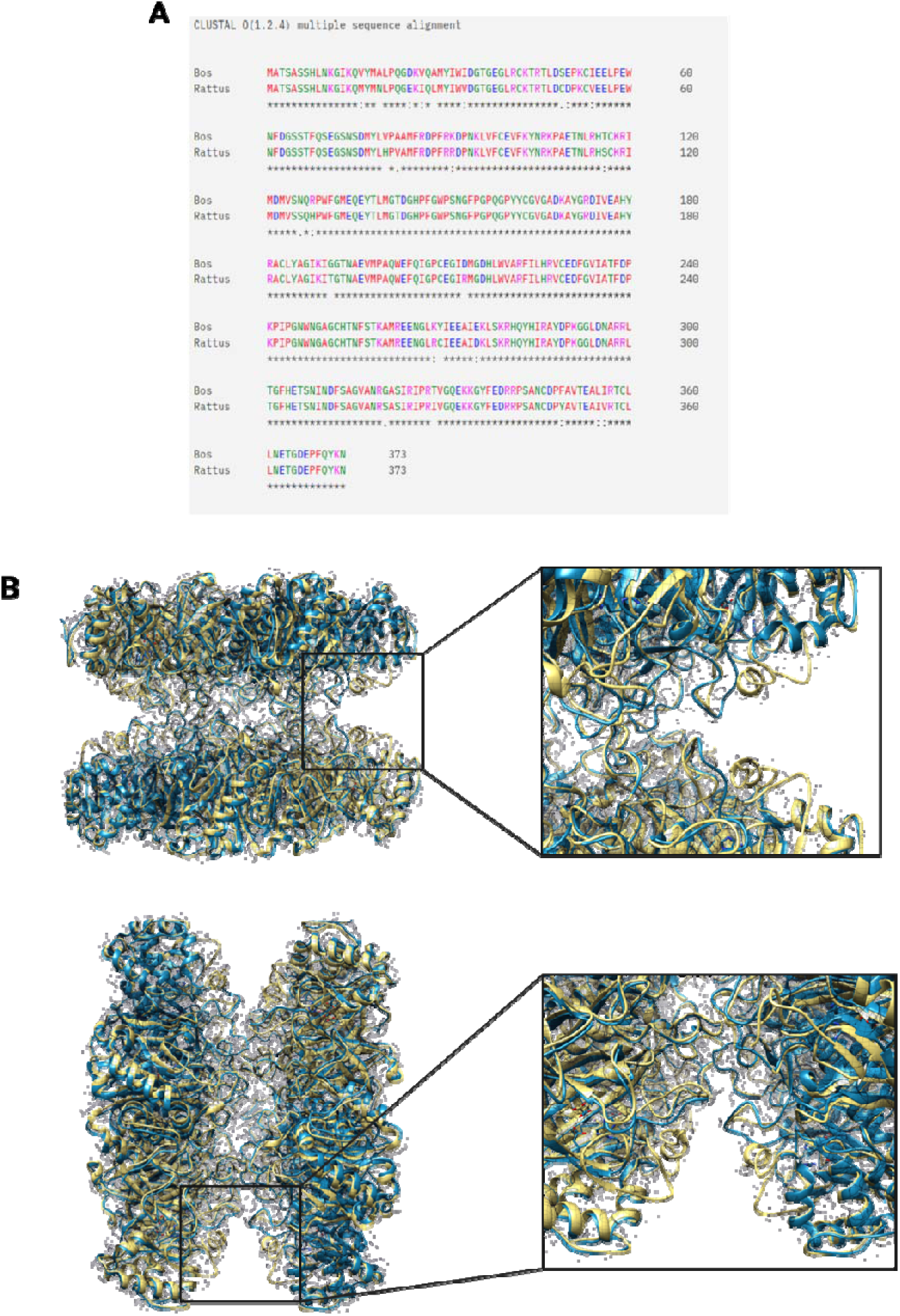
**(A)** Sequence alignment for the Bos taurus (Bovine) and Rattus norvegicus (Rat), showing the similarity at residues 291-299. **(B)** Zoomed view of the docked model of Bovine in blue and Rat in yellow into our cryo-EM density map, showing the differences at this location between two models.

**Figure S10.**
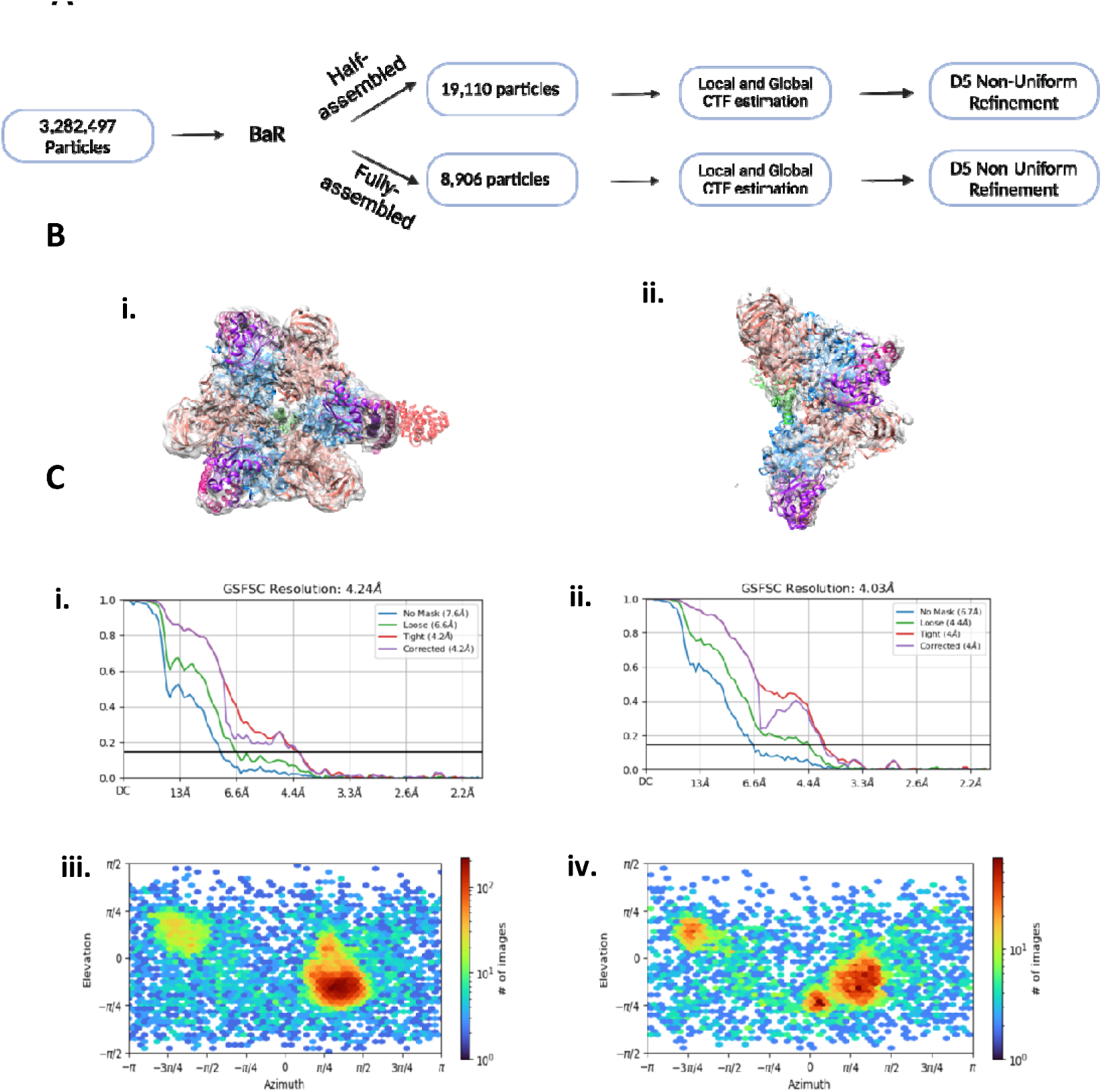
**(A)** Schematic overview of BaR analysis. **(B) i.** Side view of the reconstructed cryo-EM density map of fully assembled V_1_ V-ATPase docked into the 6vqa PDB. **ii.** Side view of the reconstructed cryo-EM density map of half-assembled V_1_ V-ATPase docked into the 6vqa PDB. **(C) i.** GS-FSC curve showing final resolution at 4.24 Å for the fully assembled V_1_ V-ATPase. **ii.** GS-FSC curve showing final resolution at 4.03 Å for the half-assembled V_1_ V-ATPase. **iii.** Viewing direction distribution for the fully assembled V_1_ V-ATPase. **iv.** Viewing direction distribution for the half assembled V_1_ V-ATPase.

**Figure S11.**
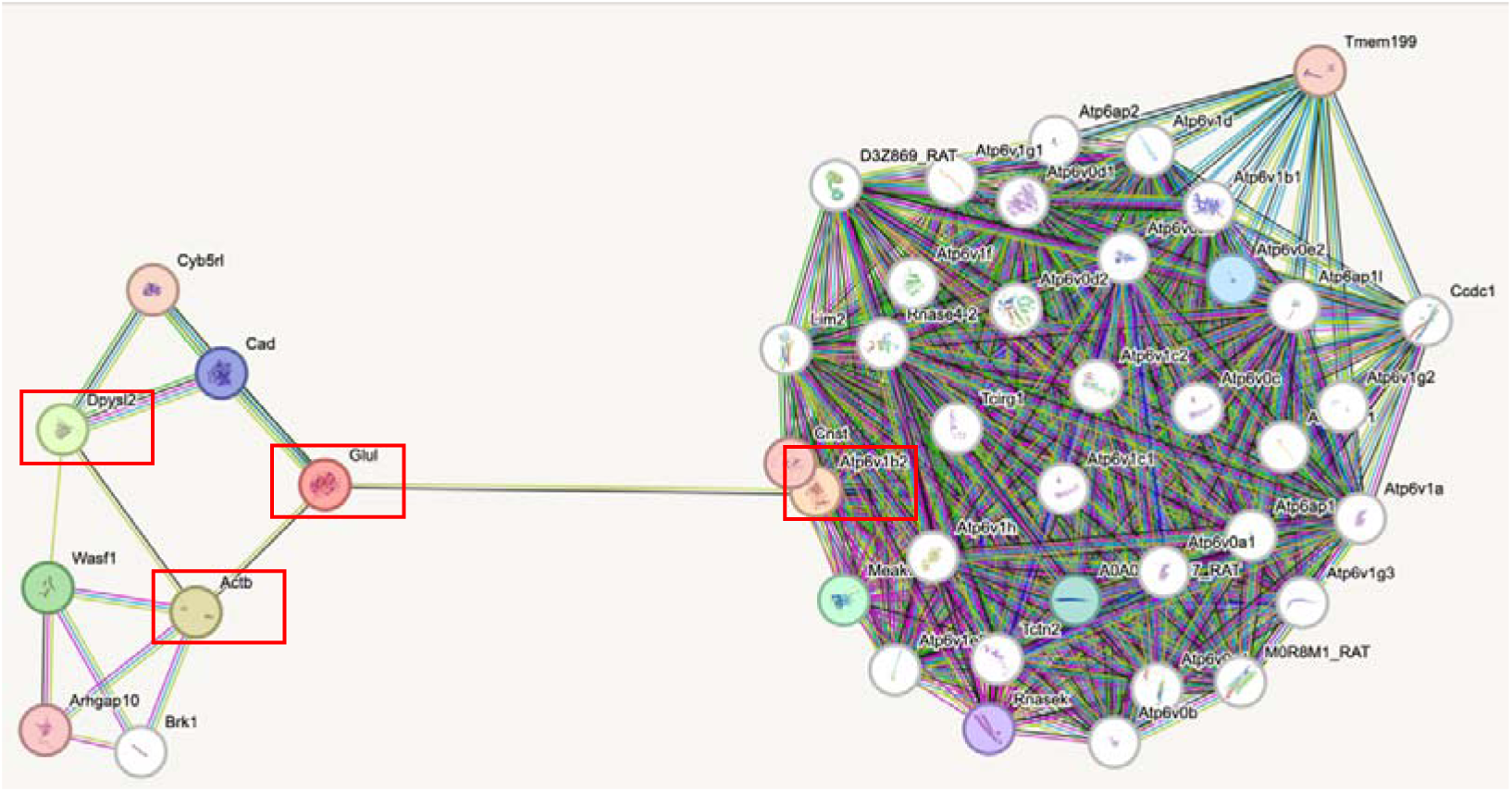
This interaction network is created using the STRING database. Lines depict interaction through different pathways. Results show all proteins identified from BaR interact through a complex network of proteins. The four enzymes, DPYSL2, GS, F-actin, and V-ATPase, are highlighted by red squares.

**Table S1.** Proteomic analysis of the phospho-modified residues in DPYSL2.

| Phosphorylated Residues | MS Dataset |
| --- | --- |
| 74 | C |
| 57 | C |
| 62 | C57 |
| 84 | C57 |
| 241 | C |
| 431 | C |
| 431 | C |
| 509 | C |
| 509 | C57 |
| 517,521 | C57 |
| 521 | C |
| 521 | C |
| 522 | C57 |
| 521 | C57 |
| 521 | C57 |

**Table S2.** Cryo-EM data collection and refinement statistics.

| **Data Collection and Processing** | **DPYSL2_D2** | **DPYSL2_C1** | **Glutamine Synthetase** | **Actin filament** | **V1 V-ATPase** |
| --- | --- | --- | --- | --- | --- |
| Microscope | Titan Krios G3 | Titan Krios G4 | Titan Krios G5 | Titan Krios G6 | Titan Krios G7 |
| Camera | FEI Falcon 4i | FEI Falcon 4i | FEI Falcon 4i | FEI Falcon 4i | FEI Falcon 4i |
| Voltage | 300 kV | 301 kV | 302 kV | 303 kV | 304 kV |
| Magnification | 75,000× | 75,000× | 75,000× | 75,000× | 75,000× |
| Pixel size | 1.03 Å | 1.03 Å | 1.03 Å | 1.03 Å | 1.03 Å |
| Exposure | 40 electrons/Å<sup>2</sup> | 40 electrons/Å<sup>2</sup> | 40 electrons/Å<sup>2</sup> | 41 electrons/Å<sup>5</sup> | 40 electrons/Å<sup>6</sup> |
| Number of Movies | 11,025 | 11,025 | 11,025 | 8,628 | 11,025 |
| Number of Frames per Movie | 30 | 30 | 30 | 30 | 30 |
| Defocus Range | 1.2-2.5μm | 1.2-2.5μm | 1.2-2.5μm | 1.2-2.5μm | 1.2-2.5μm |
| Symmetry Applied | D2 | C1 | D5 | helical | C1 |
| Initial particle images (no.) | 3,282,497 | 3,282,497 | 3,282,497 | 280,316 | 3,282,497 |
| Final particle images (no.) | 389,444 | 389,444 | 12,098 | 24,454 | 8,906 |
| Map resolution (FSC = 0.143) | 2.73 Å | 2.93 Å | 3.12 Å | 3.4 Å | 4.24 Å |
| B-Factor applied | -80.0 Å<sup>2</sup> | -80.0 Å<sup>2</sup> | -45.0 Å<sup>2</sup> | -45.0 Å<sup>2</sup> | 0 |

**Table S3.** Model building statistics.

| **Model Building** | **DPYSL2_D2** | **DPYSL2_C1** | **Glutamine Synthetase** | **Actin filament** |
| --- | --- | --- | --- | --- |
| Modeling Software | Coot, Phenix | Coot, Phenix | Coot, Phenix | Coot, Phenix |
| Number of nonhydrogen atoms | 14760<br>(Hydrogens: 0) | 14540<br>(Hydrogens: 44) | 28010<br>(Hydrogens: 0) | 16710<br>(Hydrogens: 0) |
| Number of Residues build | 1932 | 1,940 | 3,680 | 2,220 |
| Number of Ligand build | 0 | 1 | 20 | 0 |
| RMS (Bond) | 0.003 | 0.003 | 0.002 | 0.004 |
| RMS (Angles) | 0.532 | 0.596 | 0.482 | 0.616 |
| Ramachandaran Favored | 98.02% | 95.28% | 97.42 | 96.2 |
| Ramachandaran Allowed | 1.98% | 4.72% | 2.58 | 3.8 |
| Ramachandaran Outliers | 0.00% | 0.00% | 0.00% | 0.00% |
| C-beta outliers | 0.00% | 0.00% | 0.00% | 0.00% |
| Rotamer Outliers | 0.82% | 4.24% | 0.04% | 0.00% |
| All Atom Clashscore | 6.74 | 43.22 | 9.23 | 12.28 |
| MolProbity Score | 1.37 | 2.92 | 1.61 | 1.86 |
| Model-Map CC_mask | 0.86 | 0.73 | 0.84 | 0.84 |
| PDB ID | 9C4E | 9C86 | 9C28 | 9C21 |

